# Flexible Representations of Odour Categories in the Mouse Olfactory Bulb

**DOI:** 10.1101/2020.03.21.002006

**Authors:** Elena Kudryavitskaya, Eran Marom, David Pash, Adi Mizrahi

## Abstract

The ability to group sensory stimuli into categories is crucial for efficient interaction with a rich and ever-changing environment. In olfaction, basic features of categorical representation of odours were observed as early as in the olfactory bulb (OB). Categorical representation was described in mitral cells (MCs) as sudden transitions in responses to odours that were morphed along a continuum. However, it remains unclear to what extent such response dynamics actually reflects perceptual categories and decisions therein. Here, we tested the role of learning on category formation in the mouse OB, using *in vivo* two-photon calcium imaging and behaviour. We imaged MCs responses in naïve mice and in awake behaving mice as they learned two tasks with different classification logic. In one task, a 1-decision boundary task, animals learned to classify odour mixtures based on the dominant compound in the mixtures. As expected, categorical representation of close by odours, which was evident already in naïve animals, further increased following learning. In a second task, a multi-decision boundary task, animals learned to classify odours independent of their chemical similarity. Rather, odour discrimination was based on the meaning ascribed to them (either rewarding or not). Following the second task, odour representations by MCs reorganized according to the odour value in the new category. This functional reorganization was also reflected as a shift from predominantly excitatory odour responses to predominantly inhibitory odour responses. Our data shows that odour representations by MCs is flexible, shaped by task demands, and carry category-related information.

## INTRODUCTION

Categorical perception is a basic and widespread feature of sensory systems [1]. In simple forms of category learning, a continuous set of stimuli are divided into discrete groups such that they become separated by a sharp category/perceptual boundary [2]. In earlier studies of category learning, distinguishable objects on each side of the category boundary were shown to be treated as the same [3, 4]. In real life, however, even when we spontaneously group two items into the same category, we do not necessarily treat them as identical for all purposes [3, 5]. For example, we can learn to classify the same object into different categories depending on context.

Neural correlates of flexible categorisation have been found in the temporal, parietal, and frontal cortices in categorisation studies using visual stimuli [6–9]. Neurons in those areas show hallmarks of perceptual categorisation - greater differences in activity in response to stimuli from different categories as compared to the same category, regardless of their exact physical appearance [8]. In olfaction, the basic categorical representation of odours was observed as early as in the olfactory bulb (OB) [10, 11] and its analogue in insects - the antennal lobe [12]. In OB of fish, for example, ensembles of mitral cells (MCs) undergo sudden and global transitions during odour morphing, as proposed by attractor models that classify odour-evoked input patterns into discrete output patterns [10]. Activity patterns evoked by the inputs representing similar odours are stable over time, ensuring reliable encoding of sensory information [10]. Nevertheless, it remains unclear whether, and to what extent, such discrete network dynamics are plastic enough to adapt sensory representations in dynamic environments. For example, can distant odours be grouped together if their context is learned to be the same?

Both functional and structural considerations suggest that the OB accommodates mechanisms involved in olfactory plasticity [13, 14]. OB circuits show several experience dependant phenomena that allude to their involvement in experience dependent plasticity. At the output level (i.e. MCs), examples include experience dependent sparsening following enrichment [15], natural experience [16] or reorganization of ensemble odour representations following repeated exposure to the stimuli over days [17–19], as well as changes in odour-selectivity and changes in synchronization following odour discrimination tasks [20–22]. Moreover, a few recent studies showed that features of the task itself have a clear trace in MC activity. Following extensive discrimination learning of two very similar odours, the initially overlapping representations of the two odours became progressively decorrelated [17, 18], and reversed back to the default representation when the task demand reversed [23]. While these studies provide compelling evidence to the changing nature of the OB circuit by task demand, the computations they sub-serve focus on discrimination. Here we asked whether correlates of categorical representation change with respect to categorical logic? Specifically, we asked how do MCs representations change if animals learn to group odour stimuli in different contexts. To answer this question, we measured the change in MC representations of the exact same stimuli when animals were required to classify odours based on different contingencies. Our results indicate that representations of categories emerge dynamically by MCs in a way that fits not only task demands but also categorical logic.

## RESULTS

### Categorical representation of odour mixtures in the olfactory bulb of naïve mice

To test how odour category is represented in the OB of naïve mice we used morphing of binary odour mixtures. In this initial set of experiments naïve animals were exposed passively to the stimuli and no learning was involved. We morphed the monomolecular odours Ethyl Butyrate (EB) into Ethyl Tiglate (ET), and Ethyl Butyrate (EB) into Isoamyl-Acetate (IAA) through a series of intermediate mixtures (odour1/odour2: 100/0, 90/10, 80/20, 20/80, 10/90, 0/100). MC responses were measured by two-photon calcium imaging in awake head-fixed Tbet-cre x Ai95 mice expressing GCamp6f in MCs (Fig. 1 A,B; EB/ET mixtures : n=7 mice, N=434 MCs; EB/IAA mixtures: n=4 mice, N=186 MCs). As expected from previous work in awake mice individual MCs responses exhibited large variability in response transitions along the morphing series [10] and their temporal dynamics[15, 17, 24] both during and after odour presentation (Fig. 1C).

**Figure 1.**
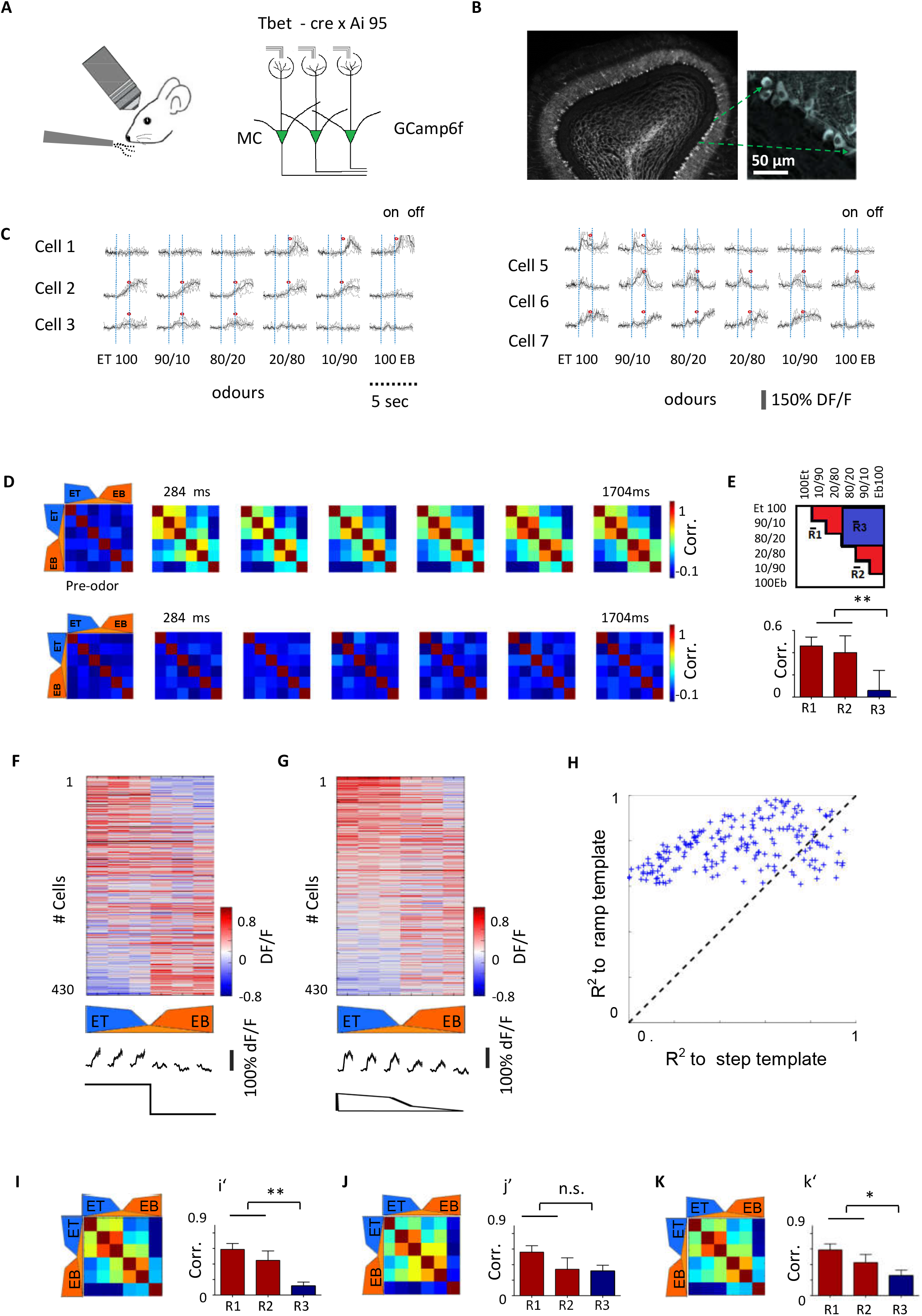
Categorical representations of binary odour mixtures by MCs. **(A)** Schematic of the imaging setup in awake naïve mice, and the OB circuit being imaged. MCs in these mice express GCamp6f. **(B)**. Coronal section of the dorsal part of the olfactory bulb taken from a Tbet-Cre x Ai95 mouse, showing that expression is limited to MC/TC. **(C)**. Calcium responses of 6 example MCs. Dashed lines indicate the odour onset and offset. Asterisk denotes a significant response. **(D)** Time series of the correlation between MC activity patterns (n=5 mice, N=230 MCs) in response to a morphed odour series of Ethyl Butyrate (EB) and Ethyl Tiglate (ET) (100/0 90/10 80/20 20/80 10/90 0/100). The first correlation matrix (left) shows correlations before stimulus onset. Top row: real data, Bottom row: same data after shuffling cell’s identity. **(E)** Top: outline of the two clusters in the matrix used to calculate within-category correlation (r1, r2, red) and between-category correlations (r3, blue). Bottom: Analysis of correlation matrix in D at 568 ms. Correlation values of the within-category clusters (R1, R2) are larger than the between-category cluster (R3; P =0.0023, Mann-Whitney U-test). (**F-G**) Matrices of calcium responses of 434 MC to a series of odours morphed from EB to ET. Responses are averaged over 1 second of odour presentation time and ranked by their correlation with a template that is shown on the bottom. (**F**) MCs responses sorted based on the step-like template that represents ideal categorizing responses of a neuron, with an abrupt transition. (**G**). Sorting based on the template that follow the ramp changes of the ratio between odours. Above each template is the average calcium response of the MCs (n= 40) with the highest match to the templates. **(H)** Scatter plot showing the relationship between the correlation coefficient values of the top ranked neurons to a ‘ramp-like’ templates (r^2^>0.5 n=200) vs their correlation to a step-like template. **(I)** Correlation matrix between MC activity patterns evoked by the morphed odours EB to ET and averaged over stimulus presentation time (same data set as in D). **i’**: Correlation values of the within-category clusters (R1, R2) and the between-category cluster (R3; P =0.01, Mann-Whitney U-test). **(J)** Same data set as in ‘I’ after elimination of 9% of neurons with the best fit to sigmoidal function (p>0.05, F test statistics). j’: R1,R2 -within category, R3-between categories. ** P =0.16, Mann-Whitney U-test). **(K)** Same data set as in ‘J’ but after after elimination of 9% of neurons with the best fit to a linear function (p>0.05, F test statistics). k’: ** P =0.03, Mann-Whitney U-test).

To test how this morphed series of binary odour mixtures are represented by the MC population we analysed the correlation among MC responses in successive time bins after stimulus onset (following an analysis described in [10]). We first combined responses of all MCs into matrices (mitral cells x stimuli), one for each 284ms time bin. We then quantified the similarity between activity patterns of MCs responses to all pairs of odour mixtures by the Pearson correlation coefficient (Fig. 1D). Morphing of one odour into another resulted in abrupt transition in the similarity of MC activity patterns at the transition from the 80/20 mixture to the 20/80 mixture (Fig. 1D-E). A second pair of odours (i.e. morphing odour EB to IAA) induced similar category-like responses (in Supl. Fig.S1 A-C). The activity patterns of MCs were separated already at a relatively early phase of the response (Fig. 1D, Suppl. Fig.S1 A,C see also [25]).

To evaluate categorization, we quantified the average correlation coefficients within and between odour category clusters in the matrices (Fig. 1E). Representation favouring odour categorisation would be expressed as high correlation within category clusters (Fig. 1E – marked as ‘R1’ and ‘R2’ shown in red) and low correlation between category clusters (Fig. 1E – ‘R3’ shown in blue). Indeed, the mean correlation of the ‘between-category’ clusters was significantly lower than the ‘within-category’ clusters for both odour pairs (Fig. 1E, bottom; Fig S1B). The mean correlation within the clusters was significantly higher than the mean correlation between clusters (ET/EB odour pair: R1 and R2 *vs.* R3, p = 0.0023; IAA/EB odour pair: p = 0.001; Mann-Whitney U-test). Thus, in the OB of naive mice, population of MCs represent odours based on the relative stimulus strength, which can be used as a natural substrate for perceptual categorization. These data are in accordance with data from multiple sensory systems, including olfaction [10, 12, 26, 27].

To evaluate the properties of MC responses responsible for mediating the transitions between odour categories, we ranked individual MCs by their response correlation to different templates. A step template represents an ideal categorizing response of a neuron, mimicking an abrupt transition (Fig. 1F, neurons on the top and bottom of the matrix are the highly correlated neurons to the step). A linear template was used to test whether neurons simply respond linearly to the odour stimuli. Since our odour concentration were designed as a ramp, a linear template is ramp-like (Fig. 1G). Linearly responding neurons for ET are shown in figure 1G. The vast majority of MCs exhibit ramp-like linear responses with different degrees of steepness to either ET or EB, and only few MCs showed step-like responses along the morphing series (Fig. 1H). To quantify neurons within each class more precisely we used an approach similar to the one described by others [10, 28, 29]. Specifically, the fluorescent responses of each MC were normalized to the range of responses of each neuron and plotted against the composition of the mixture. These curves were fit to linear and sigmoid functions (Fig. S1D). A best fit to a sigmoid function would suggest sudden transition of the response, while best fits to a linear function would suggest gradual transitions (here, a ramp response). 27% of neurons were identified as linear (P > 0.05, F statistic),a and 9% sigmoid. Because the ramp is steep separation based on linear versus step is not mutually exclusive. Indeed, some neurons that were classified as sigmoid also matched the linear function, but with lower significance. Eliminating 9% of the neurons with the higher correlation to step diminished the population transition effect (Fig. 1J,j’). Eliminating the same number of ‘linear responses’ had a smaller effect (Fig. 1 K,k’). Thus, the steepness of the response in a small subset of neurons is an important feature in OB odour classification. These data extend to the mouse similar results that were found in fish and barn owls [10, 26, 30].

Previous studies also suggested an olfactory afterimage that maintains both odour and concentration-specific information [25, 31, 32]. Interestingly, categorical representation of odours continued several seconds after stimulus offset (n=5 mice, N=230 MCs). In fact, categorical representation of odours was stronger during post odour time (Fig. S1E). These category responses are mediated by MCs that show OFF and late responses (e.g. Fig. S1F). These data indicate that categorical information is present in the odour afterimage as well [25, 31, 32].

### Sensory representation in awake vs. anesthetized mice

Previous studies have pointed to clear differences in physiological responses under awake or anesthetized imaging conditions [15, 20, 33]. We also tested to what extent the abovementioned responses in naïve mice were evident in anesthetized mice where factors like attention and state are excluded and inhibition is reduced. In anesthetized mice, MCs responses were stronger, mainly excitatory, with less trial-to-trial variability (Supl. Fig. S2A-C). Most neurons responded linearly to the morphed odour gradient (Fig. S2A-D), and step-like responses were mostly absent (Fig. S2D). Population responses in anesthetized mice also showed basic representations based on stimulus strength (Fig. S2E,G). Data from anesthetized mice were similar to the results in fish [10] where highly correlated MCs responses decorrelated over time (Fig. S2I,K). In addition, in anesthetized mice, a categorical representation in the odour “afterimage” was not evident (Fig. S2E).

### Learning of odour categorisation - same odours, different contingencies

Next, we asked whether representation of categories in the OB could develop and then change with experience. Could previously grouped stimuli be learned apart? To answer this question, we challenged mice with two go/no-go learning tasks that use the same odours but different categorical logic. 1) A 5-decision-boundary task (Fig. 2A,B), and 2) A 1-decision-boundary task (Fig. 2A,C) (see below for more details). Once learned, these tasks allowed us to assign the same odour stimuli to different category schemes. If MCs activity reflect the learning rules, odour representations would change in a way that follows categorical information.

**Figure 2.**
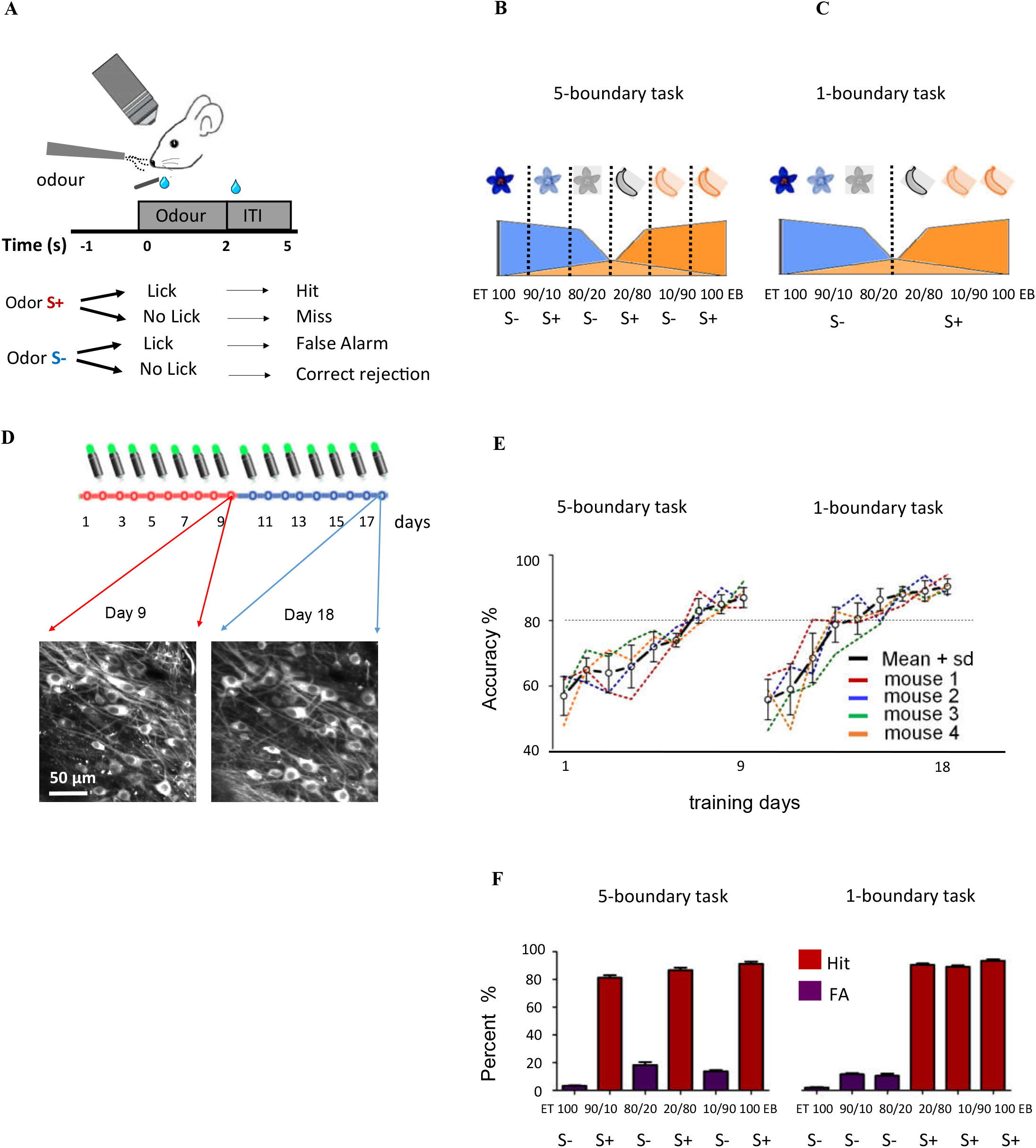
Behavioural paradigms for categorical representation. **(A)** Schematic of the go-no-go paradigm and imaging setup. The mouse is trained and imaged in a head-fixed configuration under the microscope. The mouse initiates the trial by touching the lickometer. Odour presentation begins 1 second after trial initiation. Licking following an S+ odour is rewarded with a sweet water drop. Licking following the S− odour is not rewarded nor punished. **(B)** Schematic representation of the 5 - decision boundary task with multiple discrimination boundaries intermingled along the space of binary odour mixtures (dashed lines). The S+ and S− odours include stimuli from the same and opposite sides of the natural category boundary (S+: EB100, 10/90; 80/20; S−: 20/80, 90/10, 0/100). **(C)** Schematic representation of the 1 - decision boundary task with a single discrimination boundary between ET 80/20 EB and ET 20/80EB. The S+ or S− odour value is determined according to the dominant component in the mixture (S+: ET 100/0, 90/10, 80/20; S−: 20/80, 10/90, 100 EB). **(D)** Experimental timeline. After a pre-training period in the automated homecage (see methods), mice were trained and imaged on the 5-decision boundary task for another 9 consecutive days (days marked in red). The same mice were then re-trained for another 9 consecutive days on the 1-decision boundary task (days marked in blue). Bottom: In vivo two photon photomicrographs of MCs expressing GCaMP6f from day 9 and 18 of the experiment. **(E)** Learning curves for the 5-decision boundary task (left) and the 1-decision boundary tasks (right) of individual mice under head fixation and imaging (n=4 mice). (**F**) Percentage of Hit (red bars) and FA (magenta bars) trials recorded at the last days of training (n=4 mice, mean±SEM). Left, 5 decision boundary task (9^th^ day of training). Right, 1-decision boundary task (18^th^ day of training).

In the 1-decision boundary task, we placed the decision boundary in the centre position between the odour mixtures EB 80/20 ET and EB 20/80 ET (Fig. 2C). Thus, mice were trained to assign randomly presented binary odour mixtures to S+ or S− according to the dominant component in the mixture (S+ odours: EB/ET 100/0, 90/10, 80/20; S− odours: ET/EB 100/0, 90/10, 80/20). In the 5-decision boundary task, animals had to assign three odour mixtures as S+ and three mixtures as S−, but not all based on odour similarity. S+ odours included: EB/ET 100/0, 80/20, 10/90 and S− odours included: ET/EB 100/0, 80/20, 10/90. Thus, in contrast to the 1-decision boundary task, here, the S+ and S− included stimuli from opposite sides as well as the same side of the category boundary, breaking the odor-based similarity. Such a task if learned to high proficiency (i.e. >66% correct, which is the expected level if odor-based category strategy is adopted) may force the animal to treat each item separately to break the regularity of odour-based categorisation (Fig. 2B).

Pre-training and the initial training on the 5-decision boundary task were conducted in an automated home cage setting, taking 7-9 days (Fig. S3 and see Methods). Mice were pre-trained to discriminate one odour pair, and then trained on the protocol of the 5-decision boundary task (Fig. S3). Pairs of mice were trained in the home cage for thousands of trials, until they reached high accuracy level for all stimuli (Fig. S3 B,C). We then moved the mice to the two-photon microscope and trained them in head-fixed configuration while imaging. We trained and imaged mice for ~18 days under the microscope on both tasks (Fig. 2D). Mice were trained for ~9 consecutive days on the 5-decision boundary task (Fig. 2B,E). Then, we switched the protocol to train the same mice on the 1-decision boundary task for another 9 consecutive days of training and imaging (Fig. 2C, E). Mice were able to learn both tasks at high levels of accuracy (>80%) for all stimuli, including the 10/90 and 90/10 stimuli that were switched between the tasks (Fig. 2F).

### Task-specific representation by single MCs

To measure task dependent changes in response profiles of individual MCs, we sorted their responses based on correlations between neuronal responses to task specific templates. For example, in figure 3A we show odour responses of 219 MCs recorded at days 8-9 when mice are performing the 5-decision boundary task. These neurons are sorted based on their match to a 5-decision boundary task template (a multi-step template shown on the bottom of Fig. 3A). MCs that are highly correlated (or highly anti-correlated) to the multistep template are located at the top and bottom of the matrix. Task selectivity was arbitrarily defined as having r^2^> 0.5 to the template. Unlike in naïve mice, now, only 7% (compared to 30% in naïve using the same sorting criteria) of the MCs show high correlation to a step template during the 5-descision boundary task (not shown). Thirty four percent (34%) of MCs (n=74/219) imaged at day 8-9 showed high correlations to the multi-step template (Fig. 3A,C). When we imaged the exact same MCs again at day 17-18, while mice were performing the 1-decision boundary task, those neurons were no longer correlated to the multi-step template (Fig. 3A – right). Indeed, the exact same cells that showed high correlation to the multi-step template on day 8-9 (Fig. 3C - magenta), now showed significantly lower correlation to this template (Fig. 3C - grey bars). The exact same effect (but in the opposite direction) was observed when comparing the step-like neurons while mice performed the 1-decision boundary task on day 17-18, as compared to their responses on day 8-9. Namely, at day 17-18, when mice performed the 1-decision boundary task the majority of MCs (55%) were highly correlated to a step-like template (Fig. 3B-right, 3D- grey bars). These exact same neurons responded differently one week earlier when they were imaged while the animal performed the 5-decision boundary task (Fig. 3B-left, 3D - magenta bars). Individual MCs responded based on the contingency of the odours during the specific task, and shifted their responses when task demands changed (Fig. 3E,F). Interestingly, some neurons that showed high correlation to one task template, shifted their responses to the other task template, arguing that plasticity occurs in overlapping MCs populations (Fig. 3G).

**Figure 3.**
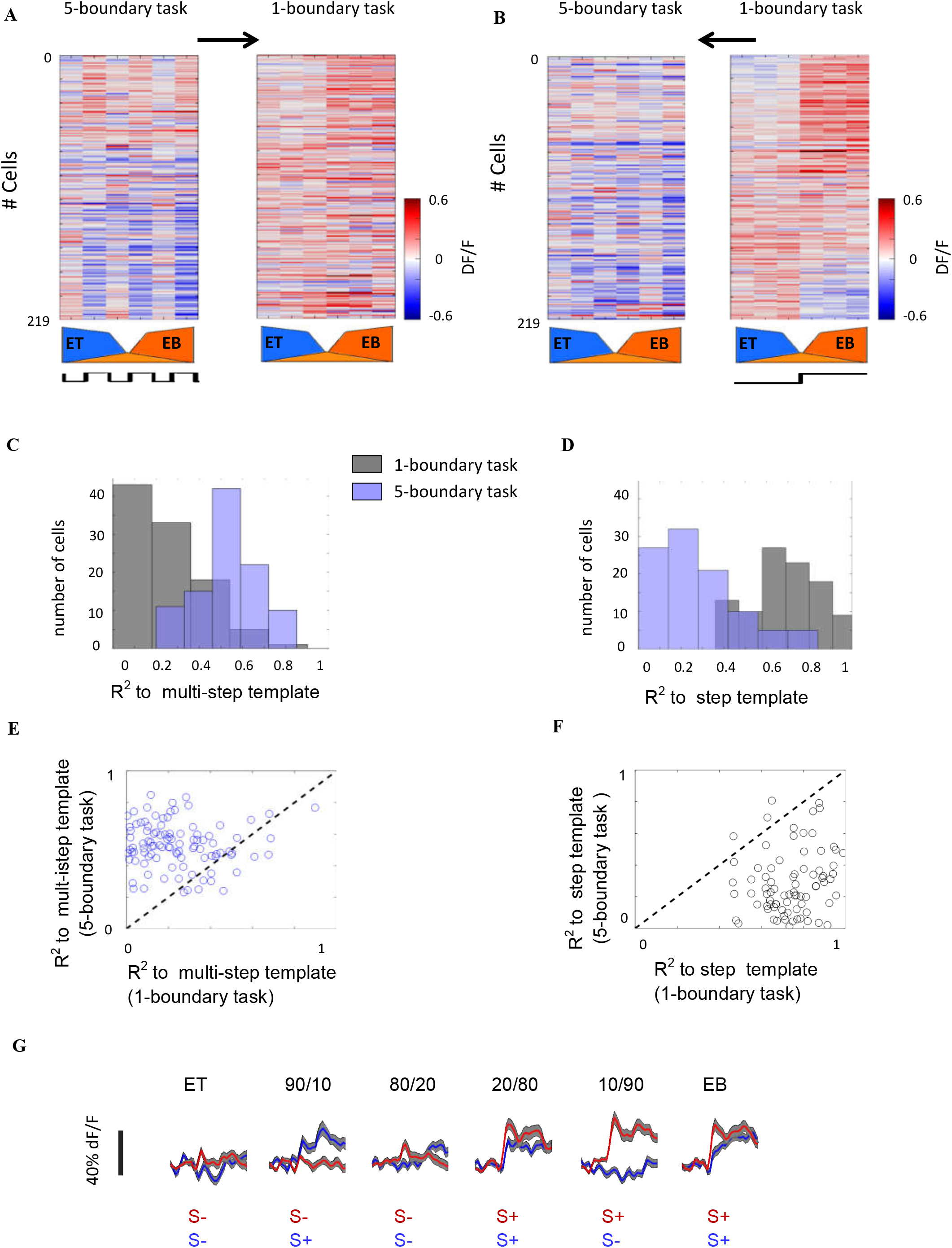
Task dependent changes of odour representations by single MCs. **(A)** Matrices of calcium responses of 219 MC to a series of morphed odours (ET to EB). Left: Responses of MCs during day 9 while the animal performs the 5-decision boundary task. DF/F of MCs responses are ranked by their correlation to a multi-step template that is shown on the bottom. The template is: −1,1,−1,1,−1,1. Responses having the best match with the template are located at the top and bottom of the matrix, respectively. Right: The exact same neurons from the matrix shown on the left but on Day 18 while the animal performs the 1 - decision boundary task. The MCs are sorted in the exact same order as in the matrix on the left and responses are to the same odours. **(B)** Right: Responses of MCs during day 18 while the animal performs the 1 - decision boundary task. MCs are ranked by their correlation to step template that is shown on the bottom. The template is: −1,−1,−1,1,1,1. Left: The exact same neurons (same sorting order and odours) from the matrix on the right but 9 days earlier on Day 9, while the animal performed the 5 - decision boundary task. **(C)** Histogram of correlation coefficients of the top ranked neurons to a ‘multi-step’ template during the 5 - decision boundary task (magenta bars), and the exact same cells 9 days later, during the 1 - decision boundary task (grey bars). **(D)** Histogram of correlation coefficients of the top ranked neurons to a ‘step-like’ template during the 1 - decision boundary task on day 18 (grey bars), and the exact same cells 9 days earlier (magenta bars). ‘C’ and ‘D’ show the 40 highest ranked MCs selected from the top and bottom of the matrices (i.e. A - 5-decision boundary task, and B - 1-decision boundary task). (**E,F**) Same data as in C and D but for individual MCs. **(G)** Average traces of responses (mean ± s.e.m) from 30 MCs selected for their high response sensitivity to a multistep template, during the 5-decision boundary task. The same neurons while the animal performs the 1-decision boundary task, 9 days later (red traces).

### Task specific representation at the MC population level

To explore the abovementioned changes of learning at the MC population level, we computed correlation matrices of population activity as mentioned above. We analysed the data with reference to different category logic as represented schematically in figure 4A,B,C (representing naïve, 5-boundary task, and 1-boundary task, respectively). Following learning of the 5-decision boundary task (Fig. 2E – corresponding to day 9), the primary category representation evident in naïve mice (Fig.4 D,d’) was completely abolished (Fig. 4E,e’; 4G). Here, the lack of the ‘naïve’ categorisation is expressed as a loss of the differences between the R1/R2 and R3 correlation clusters (Fig. 4E,e’). The correlations within clusters did not differ significantly from the mean correlation between clusters (R1 and R2 *vs.* R3; p = 0.65, Mann-Whitney U-test; Fig. 5e’). To explore the changes with respect to learning, we then analysed cell responses with respect to S+ and S− (Fig. 4B). For this analysis, we took correct trials only (see below for incorrect trials). When mice were engaged in the 5-decision boundary task (day 7-9), there was a general overall increase in similarity of responses. In addition, new task-specific boundaries were formed, which are consistent with the new behavioural demands (Fig. 4E). As expected from the single cell data, MCs ensembles represented odours based on the value associated with each odour (i.e. S+ or S−). The patterns of population responses within the S+ and S− clusters became more similar, forming two new category clusters: 1) among S+ odours (Fig. 4B, three pairs shown in red) and 2) among S− odours (Fig. 4B, three pairs shown in orange). The correlation between S+ and S− stimuli were defined as ‘between category’ (Fig. 4B, nine pairs shown in blue). The correlation values within the S+ and within the S− odours (mean±s.e.m: 0.64 ± 0.05 and 0.73 ± 0.05 for S+ and S−, respectively), were significantly higher than the mean correlation between clusters (0.55 ± 0.07; p = 0.04, Mann-Whitney U-test; Fig. 4E, e’’).

**Figure 4.**
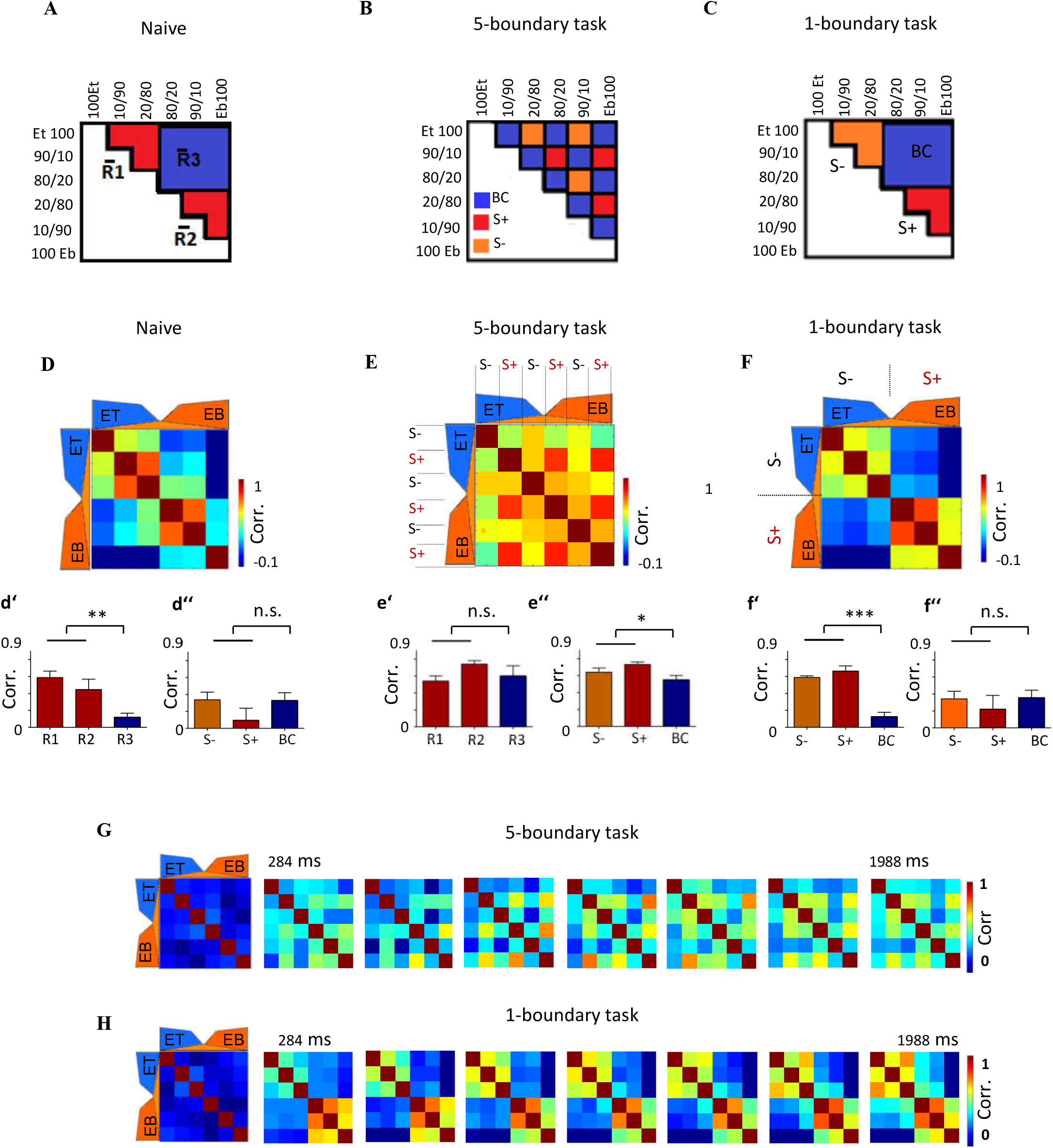
Task dependent changes of odour representations by the population of MCs. **(A), (B),(C)** Schematic inserts represent analysis of the data shown D,E, and F respectively. A and B represent analysis based on 1-decision boundary task logic. C, show data analysis based on 5-decision boundary task logic. **(D)** Pair-wise correlations matrix computed over stimulus presentation time in naïve mice passively exposed to the stimuli (Figure 1). (d’) and (d’’) analysis of the data in ‘A’ using the 1-decision boundary task logic and shown in ‘B’ using 5-decision boundary task logic, respectively. (**E**) Pair-wise correlations matrix computed during the 5-decision boundary task (correct trials only and averaged over stimulus presentation time). (e‘) Analysis of the data shown in ‘A’ using the 1-decision boundary task logic shows that the primary category representations was completely abolished. The mean correlation of all clusters was not different (r1=0.54 ± 0.06, r2=0.74 ± 0.04, r3=0.6± 0.12; p = 0.65, Mann-Whitney U-test). (e’’): Analysis of the data shown in ‘B’ using the 5-odour category logic showing that new category boundaries were formed, *de novo*. The mean correlation of the S+ and S− clusters were significantly higher than between clusters (S+ 0.73 ± 0.05; S− 0.64 ± 0.05; between clusters 0.55 ± 0.07; p = 0.04, Mann-Whitney U-test). **(F)** Pair-wise correlation matrix computed during the 1-decision boundary task (averaged over stimulus presentation time, correct trials only). Bottom (f ’)The representations of binary mixtures by the population was evident as two discrete categories. The correlation within the clusters (S+=0.56 ± 0.03, S− and S−=0.66 ± 0.03), were significantly higher than the mean correlation between clusters (0.12 ± 0.12; p = 0.0002, Mann-Whitney U-test). (f’’) analysis of the data using the 5-decision boundary category logic in ‘B’ show that 5-boundary representation was abolished following 1-decision boundary task training. The mean correlation of all clusters was not different (S−=0.34 ± 0.08, S+=0.22 ± 0.16, BC=0.35± 0.08; p = 0.65, Mann-Whitney U-test). (**G**) Time series of the correlation between MC activity patterns in response to a morphed odour series of Ethyl Butyrate (EB) and Ethyl Tiglate (ET). The first correlation matrix (left) shows correlations before stimulus onset. Data collected on day 9 during performance of the 5-decision boundary task. (**H**) Same as (**G**) but data collected on day 18th during performance of the 1-decision boundary task.

**Figure 5.**
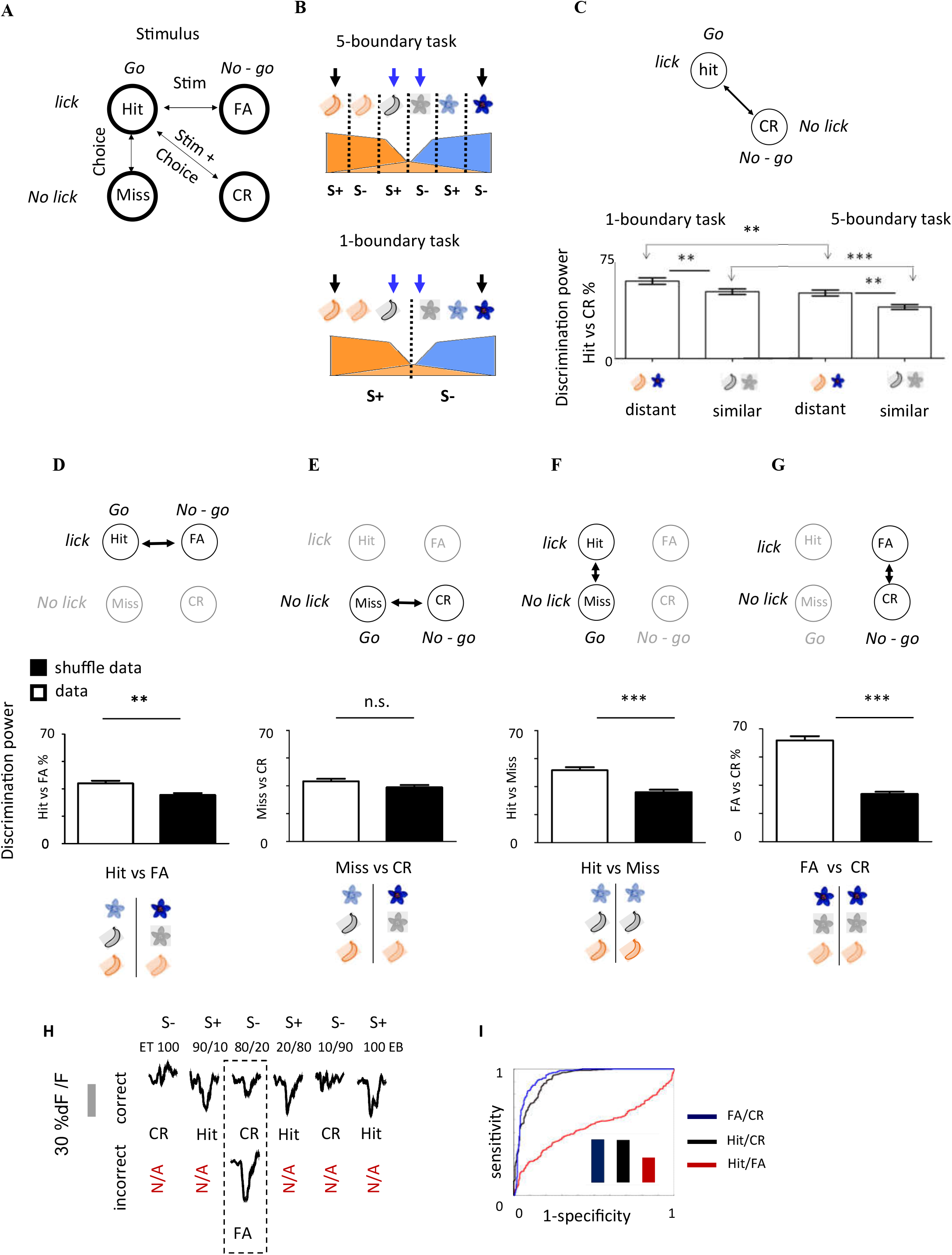
MCs responses have a strong task related component. **(A)** Schematic of the choice (lick, no lick) and stimulus (go, no-go) structure from which we derived a receiver operating curve (ROC) statistics. Calcium transients of different trials from single neurons were used to calculate “Discrimination power” from an Area Under the Curve (AUC) analysis (see Methods for details). Discrimination power refers to different pairs of trial combinations of stimuli/choices. **(B)** Schematic of the 5-decision boundary (top) and 1 - decision boundary (bottom) tasks, highlighting the two pairs of Hit vs CR trials, which we used to classify in panel ‘C’ (distant odour pair, black arrows; similar mixtures, blue arrows). **(C)** Discrimination power of MCs for Hit vs CR trials during the 5-decision boundary and 1-decision boundary tasks. Note that the odour pairs have similar contingencies in both tasks. These data show that task difficulty is reflected in MC discriminative power. Similarly, the difficulty to resolve easy and difficult odours is evident in MC activity. **(D-G)** AUC analysis (plotted as discrimination power), comparing different trial types in mice performing the 5-decision boundary task. Real data – open bars, shuffled data – black bars. (**H**) Odour-evoked calcium transients of MCs with the high selectivity to the 5-decision boundary task template (average of n=60 MCs). Top: correct trials for all odours, Bottom: incorrect trials (FA) for the 80/20 mixture. (**I**) ROC analysis and area under the curve values (bar graph, inset) classifying the MC responses in different trial types from ‘H’. Responses are different for hit vs CR and CR vs FA, but FA responses are quite similar to hits.

After animals became experts on the 1-decision boundary task, the category representation changed again, reverting back to an odour-concentration based category (Fig. 4C,F,H). The correlation within the clusters of similar odours (mean ±s.e.m.: 0.56 ± 0.03 and 0.66 ± 0.03 for R1(S−) and R2 (S+), respectively), was significantly higher than the mean correlation between clusters (0.12 ± 0.12; p = 0.0002, Mann-Whitney U-test, Fig. 4F,f’), and the 5-boundary task representation was abolished (Fig. 4F, f’’; mean ±s.e.m.: 0.34 ± 0.08 and 0.22 ± 0.16 for (S−) and (S+), respectively; mean correlation between clusters 0.35 ± 0.08; p = 0.8 Mann-Whitney U-test).

Compared to naïve mice, categorical representation became significantly higher after learning the 1-boundary task. The correlation within the S+ and S− clusters in the expert mice was significantly higher than the R1 and R2 correlation in naïve mice, when we compared the sequences shown in figure 1D to those shown in figure 4H (one-way ANOVA with Tukey’s post hoc analysis, F(5,78) = 122), p < 0.0001). No difference was found between R3 in naïve mice and the BC in trained mice. After learning, response correlations to S+ odours were higher than to S− in both tasks (Fig. 4E-e’; Fig. F-f’). Notably, the plasticity from one categorical representation to the next developed slowly over the days, and categorical representations were correlated with behavioural performance (Fig. S4). The category-like representation of task contingencies by MCs was evident only in correct trials (Fig. S5).

### MCs carry information about the task and the animal’s choice

Interpretation of odour representations in awake behaving mice is complicated by the fact that animals are engaged in the task itself. Thus, neuronal responses may also represent behavioural information like motor responses (e.g. licking), decision making, and choices (i.e. correct vs incorrect decision). We, therefore, used receiver operator characteristics (ROC) analysis of different trial outcomes to dissect the information contained in MC responses about the contribution of the task, stimuli and behavioural choices (Fig. 5A).

The most robust differences in information during the task is expected when comparing MC responses to Hit vs CR trials because these include differences in both odour identity (Hit/S+ vs CR/S−) and behaviour/choice (Hit/lick vs CR/no-lick) (Fig. 5A,C). We first compared between Hit vs CR trials of chemically distant versus chemically more similar odour pairs in the different tasks. In both tasks, these odour pairs had similar contingencies, and similar behavioural responses (Fig. 5B; distant odours-black arrows; similar odours - blue arrows). MCs showed higher discriminative power for the more distant odours in both tasks reflecting the easier nature of discriminating pure odours (Fig. 5C). Further, the discriminative power in MC responses was significantly larger for the 1-decision boundary task as compared to the 5-decision boundary task, possibly reflecting the more difficult nature of the latter task (Fig. 5C). These data show that task information is strongly represented in MC responses beyond the information about the chemicals themselves. These data are consistent with numerous reports that show task information in neuronal responses [17, 18, 20, 22, 23].

To tease apart the contribution of different task attributes we compared responses of MCs in specific trials while keeping either ‘choice’ or ‘stimulus’ constant (Fig. 5A). We first compared between correct and incorrect trials of different stimuli (only the 5-decision boundary task had enough incorrect trials for statistical power). Specifically, differences between Hit vs FA (Fig. 5D) and Miss vs CR (Fig. 5E) reflect (at least in part) a difference stemming from the chemical stimulus itself. As expected from the differential nature of the chemical stimuli, information in MCs could discriminate Hit vs FA (Fig. 5D). No differences were found between Miss and CR trials (Fig. 5E), perhaps given the large amount of trials when mice did not lick at the end of the session due to lack of thirst motivation and decreased attention. These results also suggest that decision and/or behaviour (i.e. lick *vs* no-lick) may entail a stronger component in the MC response profile as compared to the stimulus itself. To test this further, we compared MC responses when keeping the stimuli constant but choices different. Indeed, we observed a strong choice component in the MC responses based on whether they chose to lick or not (Fig. 5F-G).

An explicit example of the abovementioned result is shown in figure 5H depicting average traces of 12 trials from 60 MCs. In these MCs responses, selected for their high sensitivity to the 5-decision boundary task template, the responses for a single stimulus (ET 80/20) is different in correct (CR) and incorrect trials (FA). The difference between FA and CR trials were as high as the differences between Hit and CR trials (Fig. 5I). The responses to Hit and FA trials were rather similar, despite being chemically different (Fig. 5I). Thus, a particularly strong component in MC responses of behaving mice is related to the outcome of the trial - the stimulus value and animal’s choice - rather than the chemical stimulus *per se*.

### Reorganization of inhibitory and excitatory responses following learning

In the awake state, MCs show both excitatory and inhibitory calcium responses. The general profile of excitatory *vs.* inhibitory responses by MCs changed with learning and task demands. In naive animals, most responsive cells (71%) responded by excitation to the odours (Fig. 6A,B). Following the learning of the 5-decision boundary task, the ratio of excitatory/inhibitory responses reversed. After learning, the majority (71.4%) of neurons now responded by inhibitory calcium transients to the odours (Fig. 6C,E). The ratio of inhibitory *vs.* excitatory responses reverted back to normal after retraining the mice on the 1-decision boundary task. Specifically, 73.3% of responsive neurons were again excitatory on day 18 (Fig. 6D,E).

**Figure 6.**
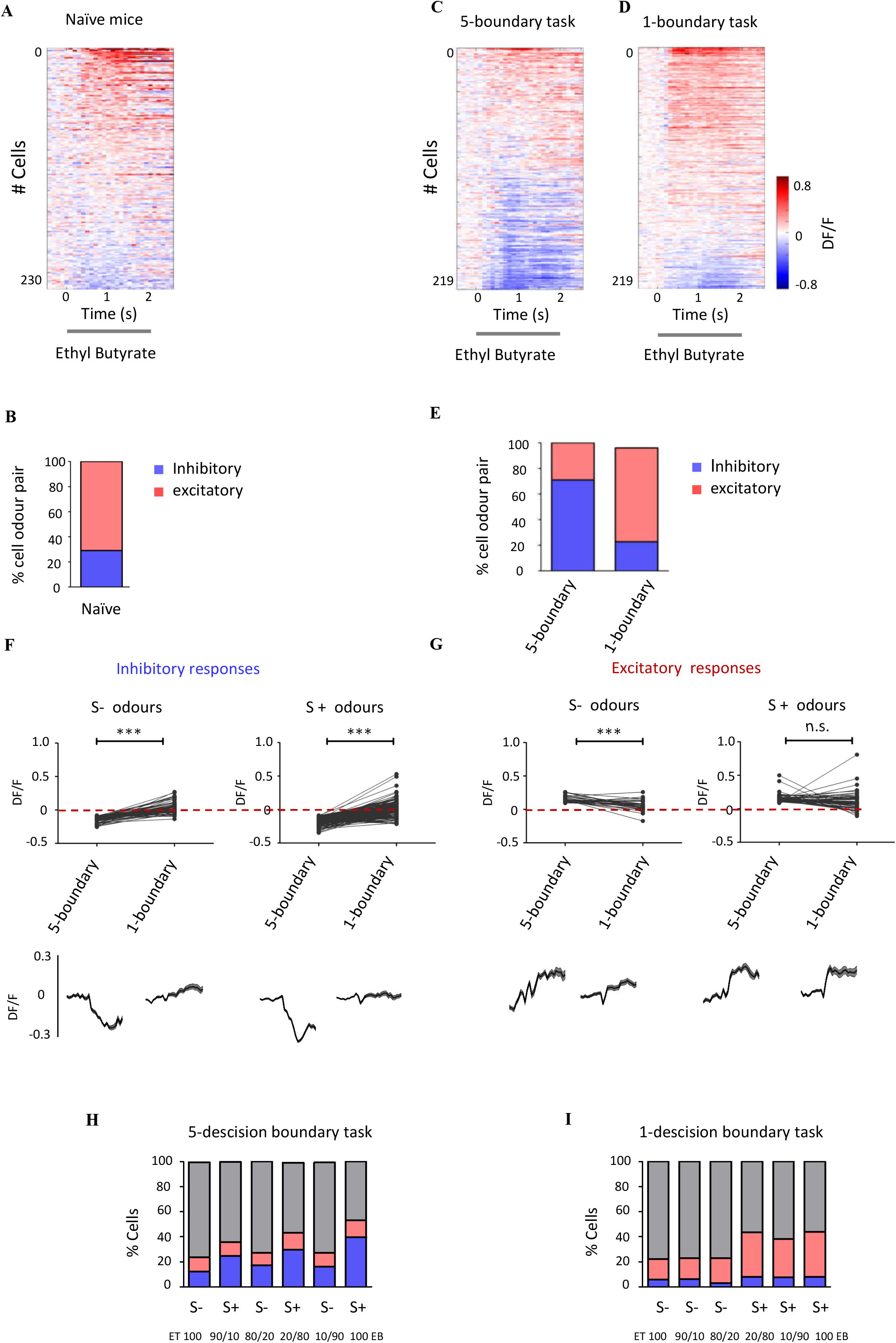
Reorganization of inhibitory and excitatory calcium responses after learning and across tasks. (**A**) Calcium responses of MCs in naïve mice passively exposed to EB. Responses are sorted by peak amplitude. (**B**) Fraction of excitatory and inhibitory responses in naive mice (n=5 mice, 230 cells, 1,380 cell-odour pairs). **(C)** Calcium responses of MCs to EB at Day 9 during the 5-decision boundary task. EB is a S+ odour. (**D**) Calcium responses of MCs to EB at Day 18 during the 1-decision boundary task. EB is a S+ odour. (**E**) Fraction of excitatory and inhibitory responses in the same MCs of mice performing different tasks (n=4 mice, 219 cells, 1,314 cell-odour pairs). (**F**) Changes in pairwise peak response amplitude of individual MCs responses from the 5-decision boundary task (day 9) to the 1-decision boundary task (day 18). Response peak-amplitude were used to classify MCs as inhibitory during Day 9 (below the dotted line) or (**G**) Excitatory responses (above the dotted line at the 5-boundary task) decrease only for S− odours. Left: Responses to S− odours (ET100 and et 80/20EB; left), Right: responses to S+ odours (EB100 and EB 80/20ET, right). Below each graph in F and G are the odour responses of all the individual cells pooled from all mice. (**H,I**) Fraction of excitatory and inhibitory responses in the same MCs calculated for each odour stimuli in mice performing different tasks (n=4 mice, 219 cells). H. 5-decision boundary task (day 9). I. 1-decision boundary task (day 18).

The main change from the 5-decision boundary to the 1-decision boundary task was observed in responses that were inhibitory. Specifically, individual neuronal responses changed sign from inhibitory to excitatory (Fig. 6F). Responses that were excitatory in the 5-boundary task showed weaker changes with learning, and even these were restricted to the S− odours (Fig. 6G, left). Regardless of the response sign (excitatory or inhibitory), the responses to S+ were always more pronounced in amplitude and number in both tasks (Fig.6 H,I). Response amplitudes of MCs following learning were lower compared to the naïve state, consistent with numerous studies [15, 17, 18].

## DISCUSSION

We imaged odour-evoked activity from MCs in response to morphed stimuli in naïve and behaving mice as they learned two tasks with different classification logic. In naïve animals, MCs reliably classified morphed odour stimuli based on the relative odour strength in the mixture. In behaving mice, when a different value was arbitrarily assigned to the same odours, odour responses by MCs changed according to the contingency learned. Our data shows that odour representations by MCs is flexibly shaped by learning and task demands, and carries category-related information.

### Categorical representation of morphed odours in the OB of naive mice

Discrete dynamics that emerge in brain networks have been proposed to underlie category formation [27, 34]. In olfaction, such dynamics have been observed as early as in the OB. For example, in the OB of fish, ensembles of mitral cells (MC) undergo sudden and global transitions when presented with morphed odour stimuli, as proposed by attractor models [10]. Similarly, in our study, MCs ensembles show category-like responses to odour stimuli without any prior odour category learning (Fig. 1; Fig. S1). Similar to the study in fish, transitions from one representation to another were mediated by a small number of neurons. Only a small number of MCs changed their activity abruptly, responding to one or another side of the category boundary (Fig. 1). Our data, thus, supports the notion of an attractor network dominated by a few neurons favouring categorisation.

In contrast to data in fish, in awake mice we did not observe population decorrelation over slow time scales. It has already been described that M/T activity patterns in the mouse OB decorrelate within the first breath [25]. Decorrelation was suggested to be a mechanism sub-serving fast odour discrimination. But categorical representation, which increased progressively with time, may serve computations on a longer time scale. One possibility is the coexistence of several read-out streams. For example, a fast code for early odour discrimination and a slower one that conveys information such as categories [11, 35].

### Flexible categorisation following learning

When animals learned to classify odours by their associated meaning rather than by their chemical similarity, odour representations by MCs changed accordingly (Fig. 4-5). Such correlates of flexible categorisation have been shown for the visual modality in higher brain regions like the prefrontal cortex [6–8, 36]. In olfaction, only few studies measured brain signals in the context of odour categorisation. For example, human fMRI studies suggest that perirhinal, orbitofrontal, piriform, and insular cortices play a role in category learning [36, 37]. How could information about odour category arise in the OB, so early in the sensory hierarchy?

In comparison to other sensory regions at a similar level of the hierarchy, the OB receives massive feedback projections from cortical, subcortical [38–46] and neuromodulatory centres [47–49]. These regions may convey back to the OB information that is not primarily sensory. For example, it has been shown that neurons in orbitofrontal cortex and the amygdala respond in a context-dependent manner and can develop divergent activity between reinforced versus the unreinforced odours during olfactory discrimination tasks [50–56]. Posterior and anterior piriform cortex also show plastic neuronal changes during reversal learning [56–58]. The reciprocal anatomical connections between the amygdala, OFC, PPC and OB [44, 59, 60] could allow these valence-related categories to influence sensory processing as described here. In this respect, not all aspects of categorical representation of odours can be observed in anaesthetised animals where cognitive states such as attention and motivation are not involved (Fig. S2). These differences likely result from weaker input from top down projection and weaker local inhibitory mechanisms, both of which could contribute to categorical responses by MCs [12].

MCs responses to odours are shaped through multiple inputs of various types of interneurons, most of which are local [61–65]. Cortical feedback onto inhibitory neurons [39, 40, 66] might contribute to the context-dependent reorganization described here by supressing MCs activity of incongruent stimuli. Theoretical and experimental studies suggest that odour recognition relies on matching sensory input patterns to a previously generated stimulus template that is formed by experience [67–69]. If the correspondence between the input and the stored template is low than the non-template information can be suppressed [64]. In our experiment, when animals were engaged in the 5-decision boundary task, some stimuli no longer matched the primary sensory representation (e.g. EB90/10 and 10/90ET). Based on the abovementioned theory, since the input and template did not match - MCs were inhibited (Fig. 6). When animals re-learned the 1-decision boundary task, odour values are presumably matched again to the primary sensory representation and the amount of inhibition onto MCs reduced again. OB inhibition, therefore, may be a mechanism by which the circuit matches learned odour templates in comparison to the primary (i.e. chemical) representation of odours that predominates MC response (as can be seen in anesthetized mice).

Could the responses that we measure be due to reward per se? Several results argue against this possibility. First, the plasticity from one categorical representation to the next did not change abruptly with the changing reward pattern but rather developed slowly over the training days along with performance (Fig. S4). Second, MCs response profiles changed dramatically with the nature of the task. Most of the responses shifted to inhibition after animals learned the 5-descision boundary task. This transition cannot be explained by lick activity alone. Third, the water reward was given after the stimulus ended during the inter-trial interval (Fig. 2A). Lastly, MCs response did not increase after the stimulus ended, a time when animals were rewarded with water and licked more intensively. While we cannot rule out completely that individual cells do follow the motor activity, a more likely explanation is that responses follow odour classification and the animal’s choice.

Dynamic changes in the OB following learning were reported by other studies as well [17, 18, 23, 70]. Interestingly, when animals learned to discriminate between two very similar odour mixtures, the representation of odours by MCs diverged, improving discriminability [17, 18]. However, when the task demand was different and animals were forced to group the same odour mixtures together, the representations of the exact same odours converged again [23]. Our data extends these observations, showing that neural signatures in the OB correlate with even more abstract representations that encode the behavioural relevance of stimuli. Conceptually similar observations were obtained in rats performing a discrimination task with different reinforcers [20].

### Categorical representation in an odour afterimage in trained and naïve mice

Stimulus aftereffects have been studied in vision [71–75], audition [76, 77], touch [78–80], as well as in olfaction [17, 31, 32, 68]. We extended these findings by examining how categorical information is represented in post odour traces. We show that categorical information remains in MC activity even when odour stimuli ceased (Fig. S1 E,F). Although the afterimage signals may be generated in the OB, cortical processes likely modulate them [32–35]. Interestingly, categorical representation in an afterimage were not observed in anesthetised mice (Fig. S2I). Following learning, the after effects by MCs were highly informative about the task (Fig. S6). Since post odour representation is likely irrelevant for immediate decision, it may play a role in other processes. For example, memory consolidation and recall of odours from memory storage may use such late information broadcasted by MCs. In humans, odour-specific ensemble patterns emerge prior to odour stimulation, possibly fuelling reliable prediction for subsequent behavioural performance [81]. Thus, the traces observed after odour cession may help the olfactory system to consolidate predictions about the upcoming stimulus. As we show here, the activity in the OB is more than a simple relay system. An odour hitting the epithelial surface, induces a rich pattern of responses that go far and beyond the chemical nature of the stimulus. Surprisingly, only one synapse away from the nose.

## Methods

### Animals

All experiments were approved by the Hebrew University Ethics Committee on Animal Care and Use. In all experiments we used adult male mice (6 -12 weeks old) from a cross between the Tbet-Cre line [82] and the Rosa26-GCaMP6f line [83]. In the adult OB, Mice of this crossing express the genetically encoded calcium indicator GCaMP6f specifically in MCs/ TCs. MCs could further be identified by their morphology and depth position within the OB (approximately 280microns under the pia matter).

### Chronic window implantation over the olfactory bulb (OB)

For the time-lapse calcium imaging of mitral cells (MCs) we used a cranial window preparation, which was described in detail in previous work [84, 85]. Briefly, mice were anesthetized with an intraperitoneal injection of ketamine / medetomidine and carprofen analgesia. A small, piece of glass (~1mm diameter) was cut from a cover slip using a diamond knife. The skull overlying one of the OBs was carefully removed using a micro-driller, leaving the dura intact. Then, a custom-made rectangular glass coverslip (No. 1) was positioned over the opening. The window was then sealed in place using histoacryl (TissueSeal) and dental cement. A 0.1 g metal bar was glued to the skull for repositioning the animal’s head under the microscope in consecutive imaging sessions [86]. Non-steroidal anti-inflammatory treatments (Carprofen, i.p.) were given 3 consecutive days post-operatively. All behavioural sessions began more than 3 weeks after the surgery.

### In vivo two-photon calcium imaging

Imaging was conducted with an Ultima two-photon microscope from Prairie Technologies (Middleton, WI, USA). GCaMP6f was excited at 950 nm (using the Deep See femtosecond laser from Spectra-physics) and images (420 × 190 pixels) were acquired at 7 Hz, using a 16x objective (NIKON) at depths 280–400 microns below the surface of the OB. Field of views were re-identified by the surface blood vessel pattern using an epifluorescence lamp and the x, y, z stage co-ordinate. Imaging cycles were triggered by the odour delivery system. In experiments when mice were exposed to the odours passively animals were habituated under the microscope in the head-fixed position before the imaging session. Each mouse was habituated once a day for about 15 min for several consecutive days until animals showed no obvious signs of stress. For the imaging experiments during behaviour, mice were habituated to the imaging conditions during the initial phase of training.

### Data acquisition and pre-processing

Data in this work were obtained from awake, head-fixed mice imaged passively or engaged in the tasks, as well as anesthetized mice. Imaging in awake animals while training took place after approximately 3-4 days after switching from home cage pre-training setting (Fig. S2).

During these days animals were habituated to the imaging procedure and training set up. Each session lasted, on average, for about 1-1.5h depending on the level of motivation. The laser power was adjust to minimize photo bleaching. Data were acquired at the last days of training (day 7-9) when accuracy level exceed learning threshold of 80%. Trials when animals showed no motivation to lick as well as when imaging conditions were poor or strong movements were evident, were excluded from the data analysis.

The odour presentation protocol included 6 odour stimuli - mixtures of two monomolecular odours and a blank channel with no odour inside. On the first imaging session, 1-2 fields of view were recorded for each mouse for a total of about 70-80 MCs per mouse and the same fields were monitored throughout all training sessions on both tasks. The fields were chosen in a way that MCs in each field showed responses to both odours in a mixture. Each trial was composed of 5 s pre-stimulation, 2 s odour presentation, and 5 s post stimulation for a total of 12 s. In awake animals, the number of trials varied with the motivation. In passive exposure experiments, odour presentation was repeated six times in each imaging field. Imaging data were pre-processed using ImageJ software, followed by a custom scripts written code in MATLAB (The Math Works). Image frames were corrected for infocal (XY) plane brain motion using ImageJ plugins [87]. Regions of interest (ROIs) for the analysis of calcium signals were drawn manually over all MCs cell bodies in each image plane. Fluorescence intensity was averaged across all pixels in each ROI, resulting in a time series of fluorescence intensity values for each ROI and trial. To calculate the relative change in fluorescence (ΔF/F), the baseline fluorescence F0 was defined as the mean fluorescence during the four seconds preceding stimulus onset. The same procedure was used to determine F0 in a blank trial without odour stimulation. ΔF/F traces were further low-pass filtered using loess fitting.

### Odour delivery

We used binary odour mixtures (100/0 90/10 80/20 20/80 10/90 0/100) of monomolecular odourants obtained from Sigma Aldrich. (ethyl-tiglate, ethyl-butyrate, Iso-amyl acetate). For the odour stimulus presentation, we used a nine-odour air dilution olfactometer (RP Metrix Scalable Olfactometer Module LASOM 2), as described by others [88]. Briefly, the odourants were diluted in the mineral oil to 10 ppm concentration. The final concentration in the vials was further calibrated with photoionization detector (miniPID, Aurora Scientific) until PID response amplitude was equalized for all channels. Saturated vapour was obtained by flowing nitrogen gas at flow rates of 100 ml/min through the vial with the liquid odourant. The odour streams were mixed with clean air adjusted to produce a constant final flow rate of 900 ml/min. Odours were further diluted tenfold before reaching the final valve (via a four-way Teflon valve, NResearch). In between stimuli, 1000 ml/min of a steady stream of filtered clean air flowed to the odour port continuously. During stimulus delivery, a final valve switched the odour flow to the odour port, and diverted the clean airflow to the exhaust. Odours were delivered at a flow rate of 1 L/min for 2 s. Inter-trial interval was 5 s. All flows were adjusted to minimize the pressure shock resulting from line switching stabilization after opening the final valve. The olfactometer was calibrated using a miniPID (Aurora Scientific) and odour delivery was monitored throughout experiments. Each trial included all 6 stimuli and blank (empty channel) To prevent predictability of presented stimuli odours were presented randomly within each trial.

### Calcium imaging data analysis

Data were analysed using custom code written in MATLAB (The Math works). For the time lapse analysis, we used only cells that were clearly identified in all imaging sessions. Significant responses were defined as those in which the magnitude of odour-evoked fluorescence change exceeded 3x the standard deviation of baseline fluctuations. We compared responses of the MC population to morphed odour stimuli by applying techniques used previously to study categorisation of odours [6, 10, 26]. Population response patterns were compared, by correlation analysis, across different combinations of morphed odour pairs. The responses of all mitral cells were combined into matrices (mitral cells/ stimuli), one for each time bin (284 ms). The similarity between activity patterns (column vectors) evoked by different stimuli and at different times was quantified by the Pearson correlation coefficient The correlation values (corr command in MATLAB) were organized in matrices, the rows (and columns) of which represented the different odour stimuli.

To test for stimulus categorisation, we compared two metrics: (1) the average within category correlation (WC) in response correlations for odour pairs in the same category and (2) the average between-category correlation (BC) in response correlations for odour pairs across categories. The strength of categorisation was quantified using a categorisation index that reflects the relationship between the WC and BC values. It was defined as (WC-BC)/(WC+BC). Positive values of the index indicate high within-category values and categorisation while negative values indicate low within-category values.

Hierarchal clustering of the correlation values was performed using the Matlab function linkage. Euclidean distance was measured in the N x cells-dimensional space in which each dimension represents the Ca 2 + signal of one mitral cell. Colour was scaled individually for each matrix; The Euclidian values (pdist command in MATLAB) were organized in matrices, the rows (and columns) of which represented the different odour stimuli.

We used ROC analysis [12, 89] to evaluate individual MCs responses as classifiers for task variables (S+ or S−) as well as for the expectation of animals in error trials. In the ROC analysis, a binary classifier computes a score for each input that is thresholded to assign a class (positive or negative) to the input. The fraction of positive inputs that are correctly classified yields the true positive rate (TPR) of the classifier at the given threshold, while the fraction of negative inputs that are incorrectly classified yields the false positive rate (FPR). Plotting the TPR against the FPR yields the ROC curve. The area under this curve is the AUC score which is a measure of the performance of the classifier (0 for perfect reverse classification, 0.5 for chance performance, 1 for perfect classification). Because MCs exhibit excitatory as well as inhibitory responses to make valid comparison we treated excitatory and inhibitory responses separately. In the case of population analysis, AUC value was calculated for each neuron and then transformed to “Discriminatory power”, which is the absolute value of the AUC minus 0.5, converted to percentage. Analysis was performed for 2 s of stimulus presentation time.

### Statistical analysis

We used pairwise comparisons of normally distributed datasets, as indicated by the Kolmogorov-Smirnov test statistical tests using two-tailed and parametric paired, and the 2-sample t-test.

Statistical significance of comparisons between correlation clusters were performed using a non-parametric Mann–Whitney U-test. All correlation values are Pearson product moment correlations. The error bars in the data are presented as mean ±S.E.M.

### Behavioural setup

Pre-training and Initial training was done in an automated custom made home cage setting integrated with RFID antenna, which allowed identification of mice individually. Specific training protocols have been assigned specifically to each mouse. The setting included a home cage connected through the short tunnel with the small Plexiglas chamber (10×10×10 cm). About 15 mm diameter hole through the wall of the chamber ended with a glass tube where the odour and water were delivered. The top of the tube was connected to an exhaust line and the bottom of the tube was connected to the olfactometer. Gauge stainless steel tube ending in a 3 mm diameter ball served to deliver water reinforcement through a 10 ml reservoir via a normally closed solenoid. Operation of the solenoid dispensed 10-20 μl of water. The amount of water could be controlled by setting valve opening window through the software. Each lick closed the electrical circuit for the duration of the tongue - tube contact and the junction potential between the metal tube and the water or the mouse’s saliva could be recorded (below 1 μA current). The output was converted via custom made A/D converter and the data from lick and infra-red diodes were further processed into USB-6341board (National Instruments). An infrared diode was used to detect snout insertions into the tube. An RFID antenna was positioned above water-odour port and connected to the transceiver which communicates with a computer by means of a custom written code in Matlab (Math Works) application. All olfactory discrimination experiments were performed using 15-odour air dilution olfactometer (RP Metrix Scalable Olfactometer Module LASOM 2) controlled by custom software written in Matlab (Math Works). The mice (up to 4) were living in the home cage several weeks while training and could enter chamber with the odour-water port anytime freely throughout the day (Fig. S2).

Mice initiate each trial by breaking a light beam at the sampling port opening. This opens one odour valve and a final valve (FV) after a short time interval (usually tFV = 1s). The use of the final valve ensures that odour traveling time between “odour onset” and first contact of the animal’s nose with the odour is minimized. Also, it ensures that odour presentation is strictly limited to the odour presentation time. After the release of the FV, the odour is delivered to the animal for 2 s. If the presented odour is a S+, the mouse responds with continuous licking of the lick port. If the presented odour was a S− odour (unrewarded), neither a reward nor any sort of punishment is given. After a trial is over, the final valve closes and the odour flow is redirected to the exhaust. During the entire experiment when odours are not given, clean airflow is delivered to the mouse. Trials are counted as correct if the animal licks continuously upon presentation of a S+ odour or does not lick continuously following a S− odour. A second trial cannot be initiated unless an inter-trial interval of at least 5 s has passed. Odours are presented in a randomized scheme, in equal numbers within a given block of 100 trials.

Training while two-photon imaging was adjusted to training conditions under head fixation. Under the microscope, the beam and RFID functions were disabled. The mouse initiated the trial by touching a metal tube and odour presentation began 1 second later. Correct answers were rewarded with a drop of sweet water, and wrong answers were not rewarded nor punished. The reward time was flexibly shifted depending on the details of the trainings protocol.

### Go-No-Go olfactory discrimination

Behavioural training started one month after chronic window implantation and is based on positive reinforcement (i.e. no aversive stimuli were given). Prior to the training, each mouse was implanted, under Isoflurane anaesthesia, with Radio Frequency Identification (RFID; Trovan) microchip inserted below the skin at the back of the neck between the shoulder blades and placed in the home cage, where it spend about 7-10 days for initial training. Food was provided ad libitum the water access was limited to the water-odour port. Mice could enter the port without restriction.

During habituation sessions, animals were familiarized with the training environment and learned how to reliably insert their snout into the odour port, initiate a trial and drink. This was typically achieved within 1-2 days. Animals were then trained to associate one olfactory stimulus with a water reward by discriminating between two monomolecular odours (S+ and S−). The correct response for the S + trials was at least 4 licks. A correct response for S− trials was to refrain from licking (0-3 licks were considered no lick). The reward was a single drop of water approximately 20 uL in volume. Mice learned this task within 1-2 days. After completion of the training criterion ≥90% of correct responses, the mice were confronted with protocol of the 5-decision boundary. Animals were trained to assign three odour mixtures as S+ and three as S−. S+ odours included: EB/ET 100/0, 80/20, 10/90 and S− odours included: ET/EB 100/0, 80/20, 10/90. The odour mixtures in the olfactometer vials were exchanged 1-2 times a day depending on the number of trials. To avoid adjustment to any possible sounds of the valve click, the position of the valves and the teflon tubing were changed daily. Animals were trained in groups of two in the home cage for thousands of trials, until they reached accuracy level ≥90%. Animals with the good quality of imaging window at the end of training were selected for further training under head fixation while imaging.

### Head fixed training while imaging

Few days prior to head fixed training, mice were kept under water restriction regime (total of 1 ml/day for each animal). Initially mice were habituated to the head fixation and to the imaging environment and its noises. Animals first learned to initiate the trial and lick the water delivery tube for reward under head fixation. To separate drinking behaviour from sensory activity we postponed reward time to the to 2s after odour onset. This was achieved by gradual postponing reward time in 100 ms steps until 2s after odour onset. Animals learned to responded during the defined time window in several training days and improved further as they became experts. Mice were trained on a task every day, several times for a few hours for 9 consecutive days. The odours were renewed before each session and the position of the valves was changed daily. The imaging period started after day 3-4^th^ day. Data were acquired starting from day 6^th^. At day 10, animals started training on the 1-decision boundary protocol and were trained for another 9 consecutive days while imaging. The task was to assign randomly presented binary odour mixtures to S+ or S−, according to the dominant component in the mixture (S+ odours: EB/ET 100/0, 90/10, 80/20; S− odours: ET/EB 100/0, 90/10, 80/20; Fig. 3C).

## Acknowledgements

We thank Israel Nelken for invaluable advice on the behavioural paradigm and analyses. We thank Amit Vinograd, Israel Nelken and members of the Mizrahi lab and for commenting on earlier versions of this manuscript. This work was supported by an ERC consolidators grant to A.M. (#616063), Israeli Science Foundation grant to A.M. (#224/17), German Israeli Foundation grant to A.M. (I-1479-418.13/2018).

