## Supplemental figures for "Flexible Representations of Odour Categories in the Mouse Olfactory Bulb"

#### SUPPLEMENTAL FIGURE LEGENDS

**Figure S1.** *Categorical representation by the population of MCs*

**(A)** Correlation between MC activity patterns evoked by the morphed odours IAA and EB before stimulus onset (left matrix) and at time 284ms (right matrix) (n=4mice, N=186 MCs). **(B)** Left: outline of the two clusters in the matrix used to calculate within-category correlation (r1, r2, red) and between-category correlations (r3, blue). Right: Correlation values of the within-category clusters (R1, R2) and the between-category cluster (R3) ** p=0.001**.** **(C)** Calcium MCs DF/F responses to the morphed odour mixtures (IAA/EB) (same odours as plotted in 'A'). Responses are sorted in the right matrix based on a step-like template (bottom). **(D)** Examples of responses to an odour morphing series (ET/EB) that showed a best fit to a linear (top) or sigmoid functions (bottom). **(E)** Correlation matrices of MC activity patterns evoked by morphing ET to EB along time. Matrices are shown from stimulus onset through 2.5 second after stimulus offset. The first correlation matrix (left) shows correlations before stimulus onset. (**F**) Calcium OFF responses of 4 MCs to morphed stimuli from (D).

**Figure S2 -** *Categorical representation by MCs in anaesthetised mice.* **(A-B)** Matrices of calcium responses of MCs (3 mice, 100 MC) to a series of morphed EB to ET odours. Responses are averaged over 1 second of odour presentation time and ranked by their correlation with a template that is shown on the bottom. The linear templates follow the ramp changes of the ratio between odours. 'A'' and 'B' step-like template. Traces are averaged DF/F responses of MCs (n= 15) to the six morphed odours showing a high match with the templates in A (top of matrices) and B (top of matrix). **(C)** Scatter plot showing the relationship between the correlation coefficient values of individual neurons to a 'ramp-like' template ($r2 > 0$, n280) vs their correlation to a step-like template. (**D**) Correlation between mitral cell activity patterns evoked by the morphed odours ET to EB in successive time bins in anaesthetised mice (n 3, 100 mitral cells). The first correlation matrix (left) shows correlations before stimulus onset. In anesthetized mice, initially highly correlated MCs activity patterns underwent decorrelation during the later phase of odour response. (**E**) Correlations between activity patterns shown in 'E' after shuffling of cell identity. (**F**) Correlation between mitral cell activity patterns evoked by the morphed ET to EB odours at 1,704 ms time point. The mean correlation (±s.d.) within the clusters of similar patterns ($0.88 \pm 0.03$ and $0.8 \pm 0.02$, at 1704 ms), the mean correlation between clusters was significantly lower ($0.47 \pm 0.16$; $P = 0.001$, Mann-Whitney U-test).

**Figure S3 -** *Initial training***.** (**A**) Schematic drawing of the homecage and training set up. Initial training was done in a custom-made operant conditioning room. Mice were tagged with radio frequency identification chip (RFID) for individual detection in a training box connected the home cage on one side and an olfactometer on the other (See Methods). (**B**) Discriminability during the learning of the 5-decision boundary discriminations task. d' value was calculated for each 60 trials. The average levels d' values of all mice at the end of the learning was d'=4.0±0.49(s.d). **(C**) An example ROC curve and d' of one mouse calculated from the last 200 trials of the 5-decision boundary task learning for different odour mixture comparisons ET 100/0 vs 0/100EB (black); ET 80/20EB vs ET20/80EB (blue) and ET 90/10EB and ET 10/90EB (orange).

**Figure S4 -** *Evaluation of task specific changes during learning.* (**A**) Mean d' value of all mice for every two days of training after switching from the 5-decision boundary task to the 1-decision boundary task. Plotted below the graph are correlation matrices of MCs activity corresponding to the relevant day of learning. The second Y-axis (blue) corresponds to the within category-between category distance calculated form the matrices. (**B**) The distance within categories (S- and S+) and the distance between categories (BC), over the days of learning. Right- schematic depiction of the S-, S+ and BC clusters.

**Figure S5 -***Task dependent changes are more evident in correct trials. (***A**) and (**B**) Correlation matrices between MCs population responses to odours (n = 150; averaged over 1.5 seconds after stimulus onset) in correct trials (A) or when incorrect trials were dominant (B). (**C**) Schema representing groups analysed based on the category logic of the 5-decsision boundary task. Red and orange pixels represent 'Within Category' values (WC - correlation among S+ and S- stimuli). Blue pixels are the 'Between Category' values (BC – correlation between S+ and S- stimuli). (**D**) A bar graph of the category index (CI) calculated from the correlation matrices in A and B for correct trials and trials when mice performed below threshold. CI is calculated as follows: CI= (BC/(WC+BC).

**Figure S6 –** *Category information is maintained in the odour afterimage* (**A**) Left: correlation matrix between mitral cell activity patterns evoked by the morphed odours EB to ET after odour offset in naïve awake mice passively exposed to the stimuli. MCs traces were averaged over 1.5 seconds time window in the post odor time. Middle: outline of the two clusters in the matrix used to calculate within-category correlation (r1, r2, red) and between-category correlations (r3, blue). Right: Correlation values of the within-category clusters (R1, R2) and the between-category cluster (R3; P =0.0007, Mann-Whitney U-test). (**B**) Pair-wise correlations matrix computed during the 5-decision boundary task (details same as in main figure 5B, C(bottom)). The mean correlation of the S+ and S- clusters were significantly higher than between clusters (S+ 0.54 ± 0.07; S- 0.5 ± 0.07; between clusters 0.3 ± 0.12; p = 0.01, Mann-Whitney U-test). (**C**) Same as 'B' but on day 18 during performance on the 1-decision boundary task. The mean correlation of the S+ and S- clusters were significantly higher than between clusters (S+ 0.54 ± 0.13; S- 0.84 ± 0.02; between clusters 0.27 ± 0.15; p = 0.001, Mann-Whitney U-test).

### Figure S1.

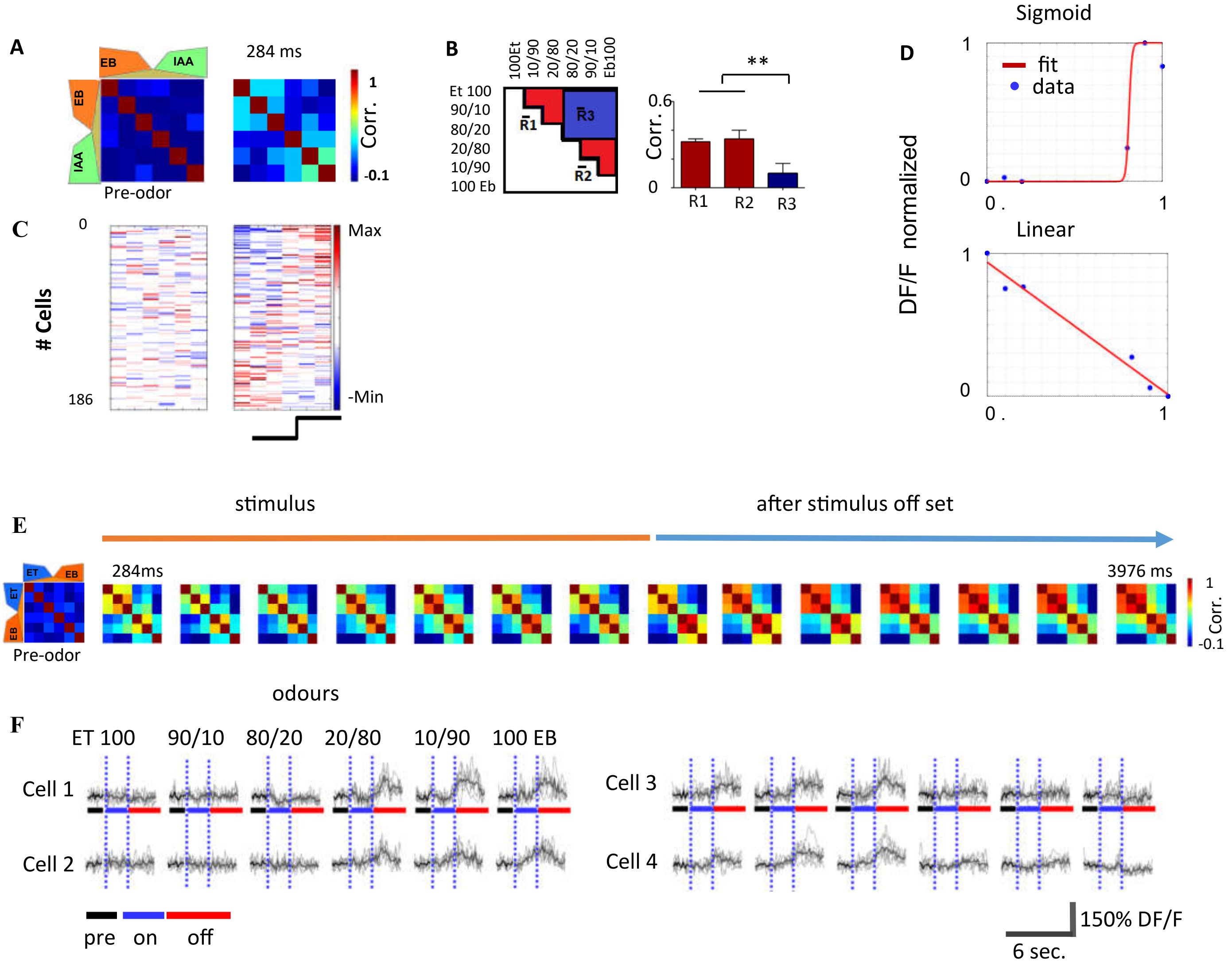
A
EB
IAA
Pre-odor
284 ms
1
Corr.
-0.1
B
100Et
10/90
20/80
80/20
90/10
Eb100
Et 100
90/10
80/20
20/80
10/90
100 Eb
R1
R2
R3
0.6
Corr.
0
**
C
0
186
### Cells
Max
-Min
D
Sigmoid
fit
data
DF/F normalized
Linear
E
stimulus
after stimulus off set
284ms
3976 ms
ET
Pre-odor
odours
F
ET 100
90/10
80/20
20/80
10/90
100 EB
Cell 1
Cell 2
Cell 3
Cell 4
pre
on
off
150% DF/F
6 sec.

### Figure S2

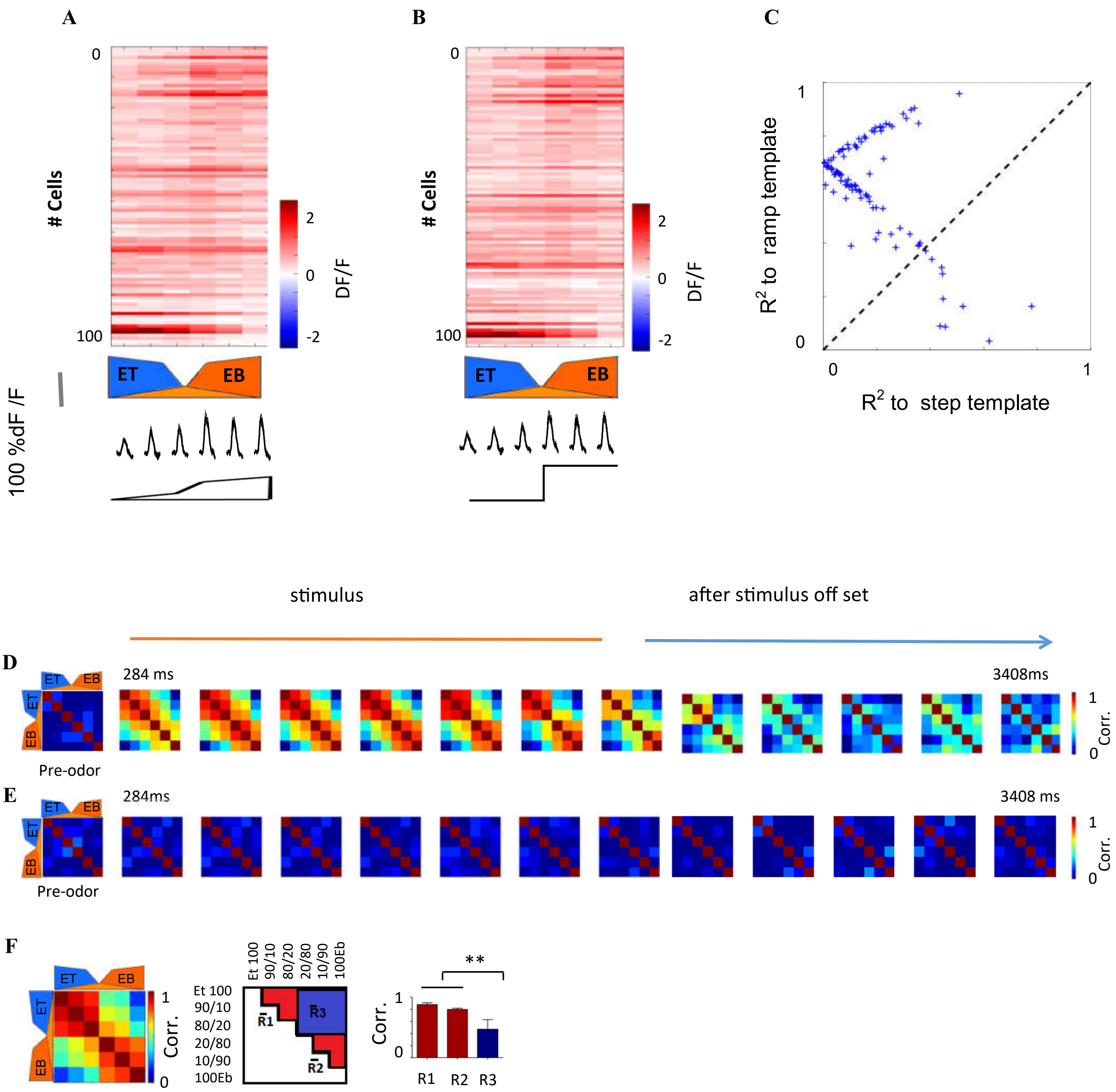

A
B
C
0
100
### Cells
2
0
-2
DF/F
ET
EB
100 %dF /F
1
R2 to ramp template
R2 to step template
stimulus
after stimulus off set
D
284 ms
3408ms
Pre-odor
Corr.
E
284ms
3408 ms
F
Et 100
90/10
80/20
20/80
10/90
100Eb
R1
R2
R3
**

### Figure S3

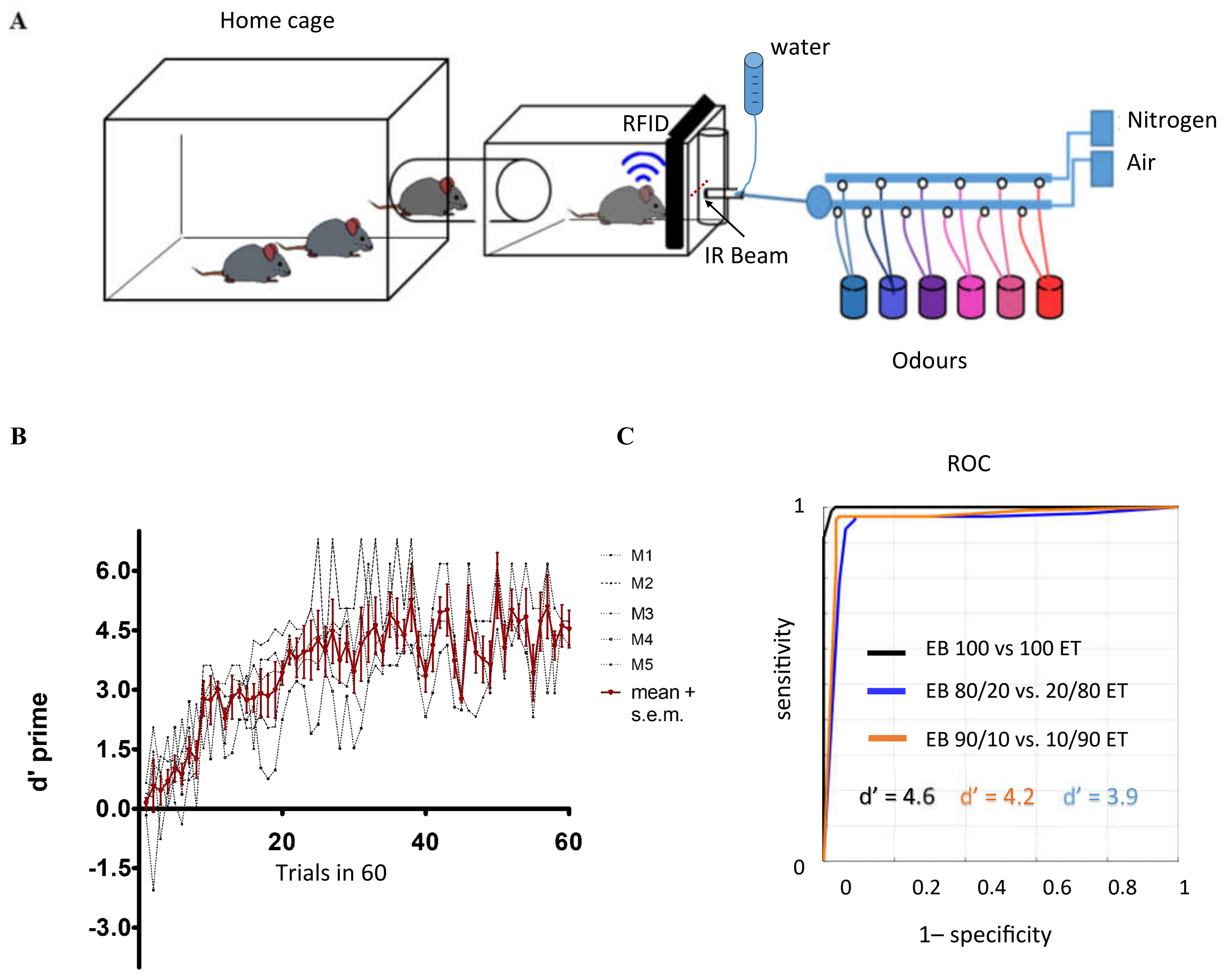
A
Home cage
water
RFID
IR Beam
Nitrogen
Air
Odours
B
d' prime
6.0
4.5
3.0
1.5
0.0
-1.5
-3.0
20
40
60
Trials in 60
M1
M2
M3
M4
M5
mean + s.e.m.
C
ROC
sensitivity
1
0
0
0.2
0.4
0.6
0.8
1
1– specificity
EB 100 vs 100 ET
EB 80/20 vs. 20/80 ET
EB 90/10 vs. 10/90 ET
d' = 4.6
d' = 4.2
d' = 3.9

### Figure S4

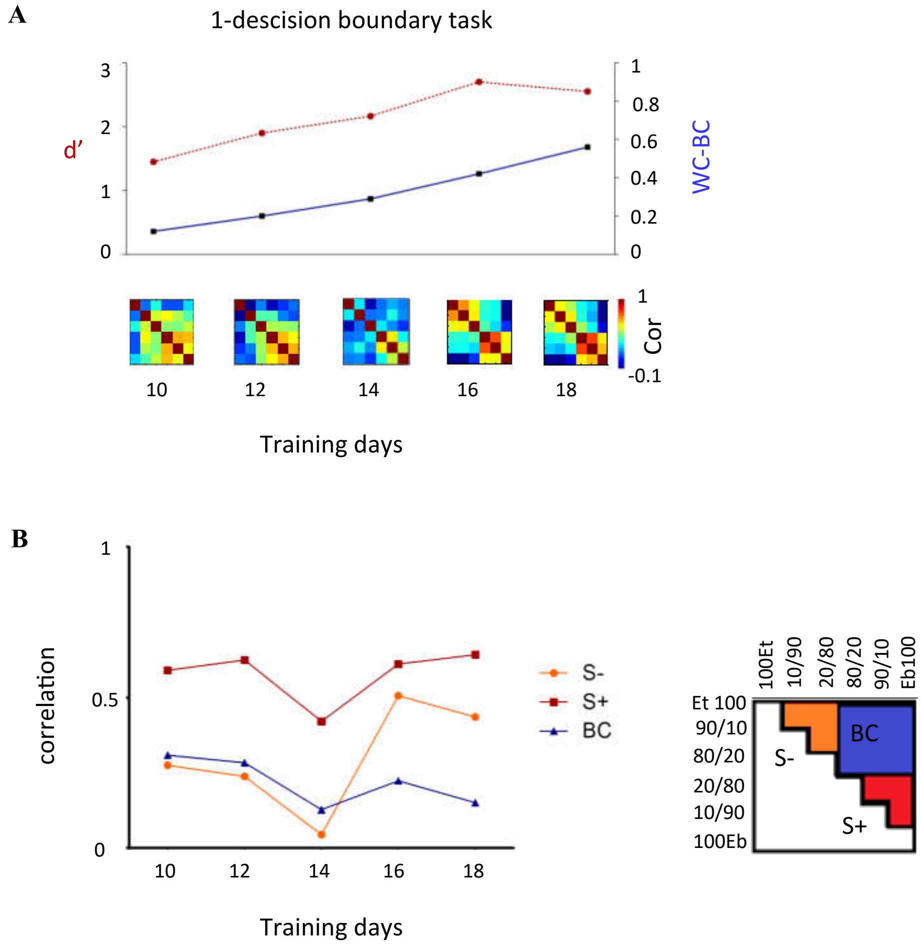

A
1-descision boundary task
d'
WC-BC
Cor
Training days
B
correlation
S-
S+
BC
Training days
100Et
10/90
20/80
80/20
90/10
Eb100
Et 100
90/10
80/20
20/80
10/90
100Eb
S-
BC
S+

### Figure S5

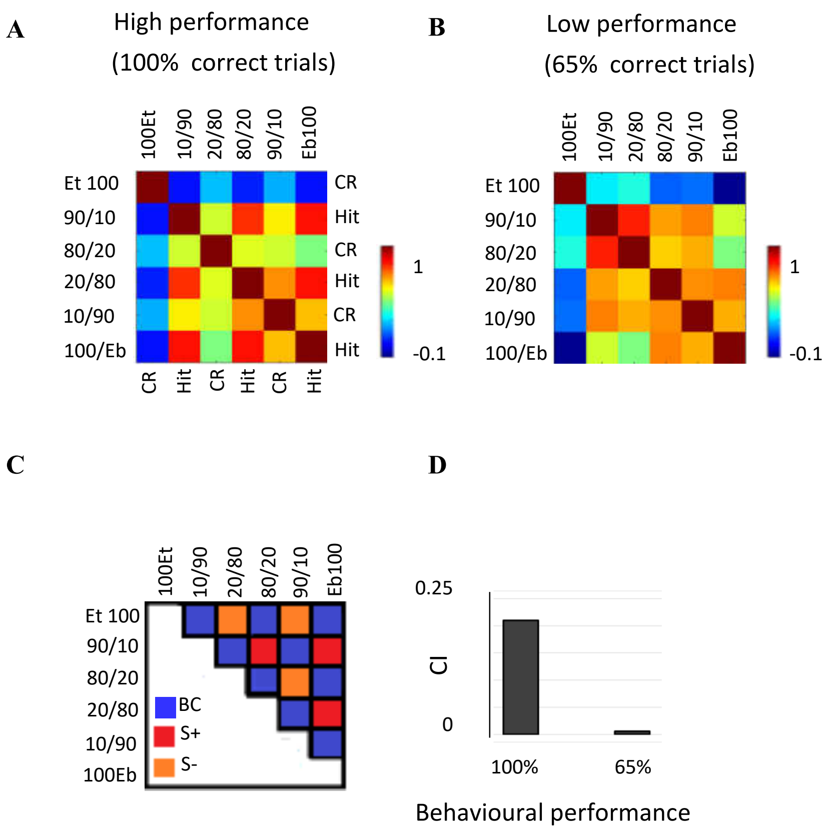
A
High performance
(100% correct trials)
100Et
10/90
20/80
80/20
90/10
Eb100
Et 100
90/10
80/20
20/80
10/90
100/Eb
CR
Hit
CR
Hit
CR
Hit
1
-0.1
CR
Hit
CR
Hit
CR
Hit
B
Low performance
(65% correct trials)
100Et
10/90
20/80
80/20
90/10
Eb100
Et 100
90/10
80/20
20/80
10/90
100/Eb
1
-0.1
C
100Et
10/90
20/80
80/20
90/10
Eb100
Et 100
90/10
80/20
20/80
10/90
100Eb
BC
S+
S-
D
0.25
CI
0
100%
65%
Behavioural performance

### Figure S6

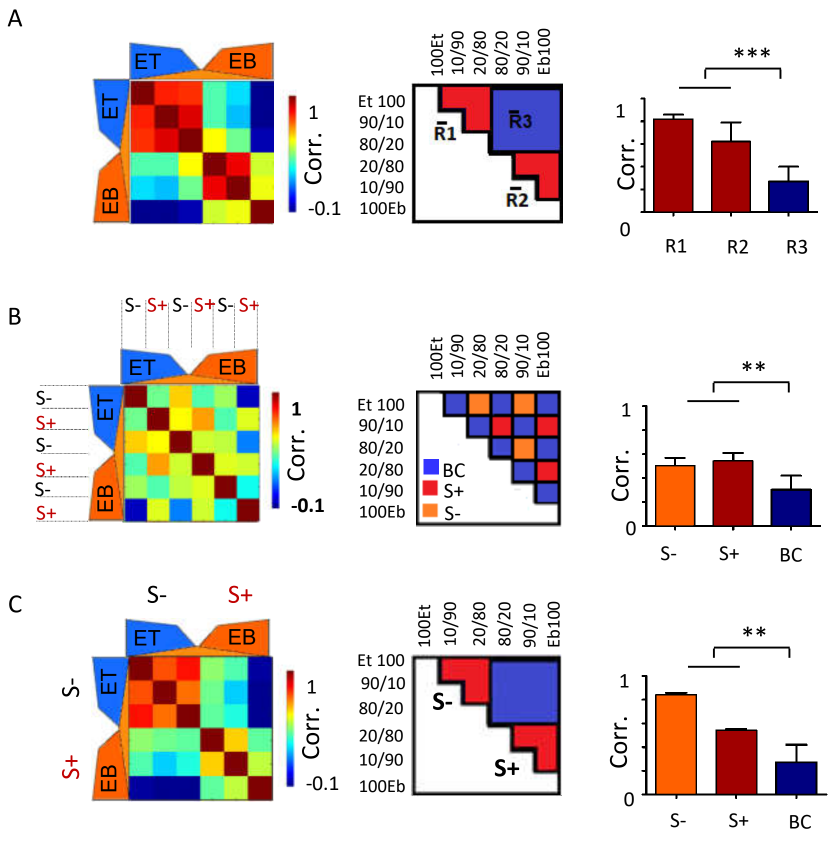
A
ET
EB
1
Corr.
-0.1
100Et
10/90
20/80
80/20
90/10
Eb100
Et 100
90/10
80/20
20/80
10/90
100Eb
R1
R2
R3
***
0
B
S- S+ S- S+ S- S+
BC
S+
S-
**
S-
S+
BC
C
S-
S+
**
